# Synaesthesia lost and found: Two cases of person- and music-colour synaesthesia

**DOI:** 10.1101/074955

**Authors:** Francesca R Farina, Kevin J Mitchell, Richard AP Roche

**Author notes:** Joint senior authors (Kevin J Mitchell, Richard AP Roche). Author email addresses: FRF; kJM, RAPR.

## Abstract

Synaesthesia is a developmental condition involving cross-communication between sensory modalities or substreams whereby an inducer (e.g. a sound) automatically evokes a concurrent percept in another modality (e.g. a colour). Whether this condition arises due to atypical structural connectivity (e.g., between normally unconnected cortical areas) or altered neurochemistry remains a central question. We report the exceptional cases of two synaesthetes – subjects AB and CD – both of whom experience coloured auras around individuals, as well as coloured perceptions in response to music. Both subjects have, in recent years, suffered a complete loss or reduction of their synaesthetic experiences, one (AB) through successive head traumas, including a lightning strike, followed by a number of medications, and the other (CD) while taking anxiolytic medications. Using semi-structured interviews and data from the Synaesthesia Battery and a colourpicker task, we characterise the phenomenological characteristics of their pre-loss synaesthesia, as well as the subsequent restoration of each subject's synaesthetic experiences (in the months post-trauma for AB, and after cessation of medication for CD). Even after years of suppression, the patterns of associations were highly consistent with those experienced pre-injury. The phenomenological experience of synaesthesia can, thus, like most conscious experiences, be modulated by pharmacologically diverse medications or head injury. However, the underlying neural substrates mediating specific synaesthetic pairings appear remarkably “hard-wired” and can persist over very long periods even under conditions that alter or completely suppress the conscious synaesthetic experience itself.

## Introduction

To date, over 60 different forms of synaesthesia have been identified (Day, 2015), while new varieties of the condition continue to be added. Recently, Ramachandran and colleagues (2013) characterised a form of emotion-evoked synaesthesia involving coloured halos around faces in a subject TK. This constituted only the eighth published report of this experience, following an initial report of a 7 year-old synaesthete in 1934 (Riggs & Karwoski, 1934), and later subjects BB (Cytowic, 1989), GW (Ward, 2004), R, F, L and M (Milán et al., 2007; 2012). Here, we report the cases of two synaesthetes – subjects AB and CD – both of whom experience coloured auras for individuals, as well as coloured perceptions in response to music. Remarkably, both subjects have, in recent years, suffered a temporary loss or reduction of their synaesthetic experiences due to head trauma and medication, respectively. We believe this constitutes the first report of the loss and subsequent return of these forms of synaesthetic experiences, and suggest that the study of such rare cases may shed light on the nature of this phenomenon.

## Results

Synaesthete AB, a 20 year-old ambidextrous (left-hand dominant) female, has had synaesthetic perceptions since childhood. Specifically, she experiences projected visual colours in response to musical notes, chords and instruments, and strong associated colours or auras in response to people. Her musical concurrents are unidirectional, and appear – typically in the centre of the visual field – as semi-transparent percepts comparable to ink or paint dropped into water. Evoked colours are temporally synchronous to the on/offset of the sounds, and are influenced by characteristics such as *pitch* (higher notes present higher in the visual field with more pastel shades; lower notes appear in lower regions with more solid colours), *volume* (louder evokes more intense colour), *type of instrumen*t (see Figure 1a for colours associated with different notes before and after loss, and Figure 1b for representations of different instruments) and *expectation* (more vivid colours for unexpected sounds). Despite being unable to read sheet music, AB can play the tin whistle (a traditional Irish flute), flute, glockenspiel, marimba and piano by ear, and cites her learning these instruments as being aided by her synaesthesia, where ‘wrong’ colours flag out of tune notes (see Figure 1c, right, for a representation of a complete song). Her person-colour associations are not projected in the visual field; rather, known individuals evoke a strong colour association in the mind's eye, with specific personality traits linked to different colours (e.g. blue people are emotional, green are loyal; see Figure 1c, left, for a list). No two people have the same colour, but colours for couples can intermingle; further, some people can have multiple colours, while a person's voice can differ from the colour of their personality. Colours are strongly influenced by emotion, and AB again states that this form of synaesthesia is beneficial in that her emotional response to a person is partly determined by their colour.

**Figure 1:**
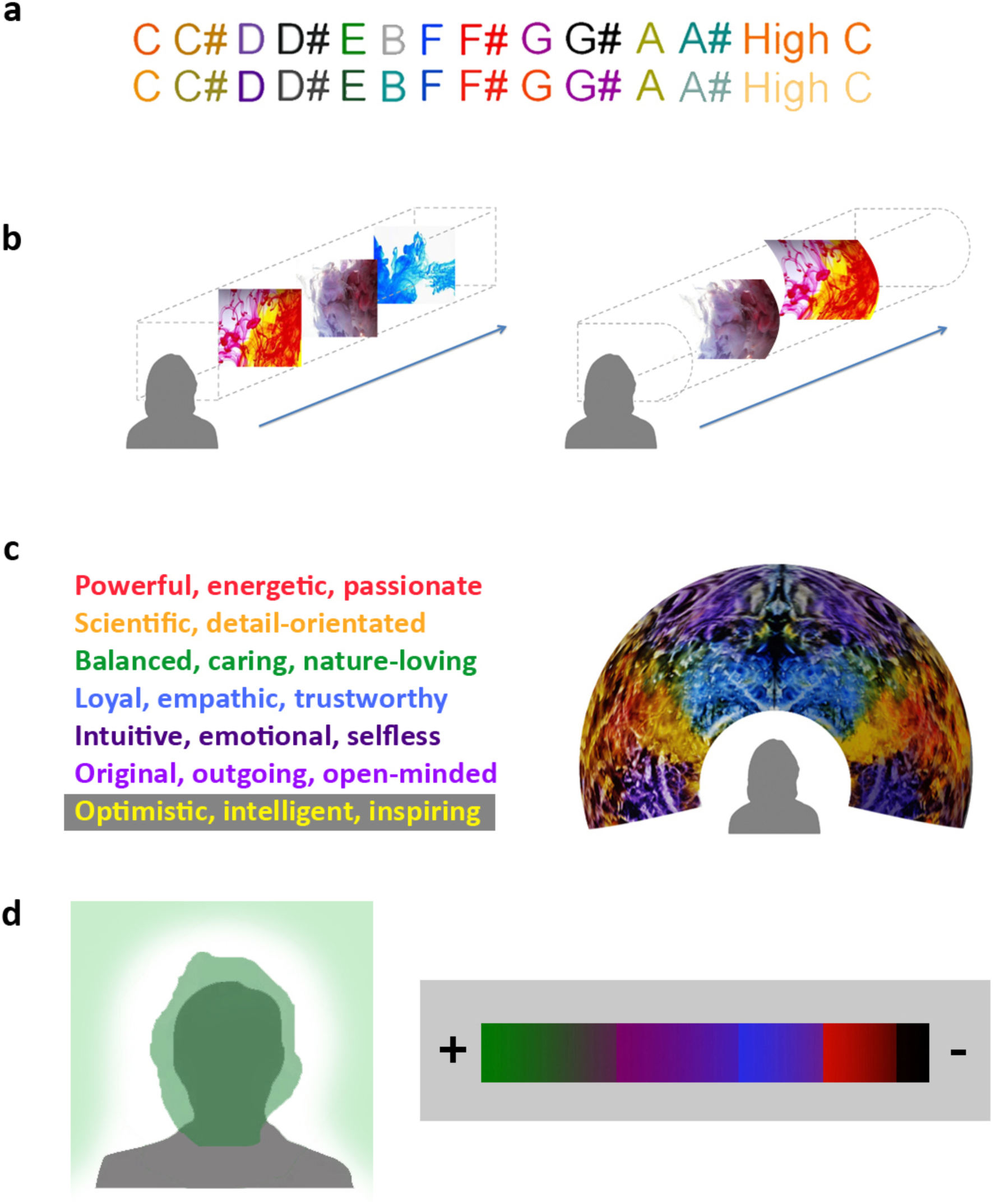
**a)** RGB-accurate colour concurrents for musical notes for AB before (upper) and after (lower) loss of synaesthetic experiences. Consistency pre- and post-loss, calculated via correlation coefficients: Red: 0.71, Green: 0.74, Blue: 0.63. Mean = 0.69. **b)** Depiction of AB's experience of music from piano (left) and flute (right) wherein colour percepts are experienced in the centre of the visual field within the confines of an imaginary box/corridor. The shape of the corridor's boundaries varies according to the instrument; e.g. for the flute, it appears “more curvy to the right side”. **c)** Left: RGB-accurate colour concurrents for AB's personality traits associated with people; Right: depiction of AB's experience of a complete song (*Flight Facilities - Stand Still*) whereby colour concurrents are described “as if projected on the inside of a dome” above her head. **d)** Left: depiction of CD's coloured auras, with independent central and peripheral concurrents; Right: RGB- accurate colour concurrents for CD's emotions, ranging from positive associations (green, purple) to ambiguous (blue, red) to negative (brown, black). Colour block sizes reflect the frequency of occurrence of each emotional association in CD's experience.

Between 2008 (age 14) and 2016, a succession of traumas caused AB to report changes in the nature of her synaesthesia (Figure 2). These initially involved two minor concussions and a short-lived suppression of her coloured concurrents while on medication (*Paramax*; paracetamol with metoclopramide hydrochloride) for migraine in 2010. On starting at university in 2012, AB was first introduced to the condition of synaesthesia, prompting her to complete the online Synesthesia Battery (Eagleman et al., 2007; see Table 1 for details). In December 2013, she contracted viral meningitis, resulting in a change in her experienced colours; they remained vivid, but were evoked by the ‘wrong’ notes and appeared ‘displaced’. After her recovery, she reports that the colours returned to normal. Three months later (March 2014), she sustained a third concussion with loss of consciousness; AB reports that following this incident, her music-evoked colours moved from the centre to the lower periphery of her visual field and appeared muted or softened, without much change to her person-colour synaesthesia. The following month (April 2014) a fourth concussion produced no loss of consciousness, but resulted in increased anxiety and emotionality, and a change in her synaesthetic experiences. These became quite intense and vibrant, and also bothersome as the colours “seemed weird” and easily led to sensory overload. In June 2014, by which time her experiences were “on their way back to normal”, AB was involved in a lightning strike (she was inside a metal cabin with her hand on the windowsill when the structure was hit by lightning), leading to a brief hospitalisation. This led to memory loss for the period of several days immediately after the lightning strike and increased anxiety afterwards, with additional changes to her synaesthetic experiences. Normal colour experiences were essentially abolished, though she did experience “an awful lot of white” due to a ringing in her ears, and later developed pins and needles, which she describes as seeing white flashes in her head. She suffered a number of panic attacks during this time and describes uncharacteristic synaesthetic experiences preceding them, including perceptions of intense, strange colours, mostly golds and silvers, which she describes as possibly “not even real colours”. Following this incident, AB was prescribed Xanax (*Alprazolam*) for one month. This caused a temporary muting of her colours for both music and people, but by July of that year they had partially returned (with high consistency – r=0.69 – to their original colours; see Figure 1a). Over this time, AB developed seizures, which led to her being prescribed Keppra (*Levetiracetam*) in December 2014. This resulted in a complete suppression of her synaesthetic colour experiences, as well as generally low mood and difficulty concentrating. After a month of increasing dosage of this medication, AB was removed from it due to the severe side effects.

**Figure 2:**
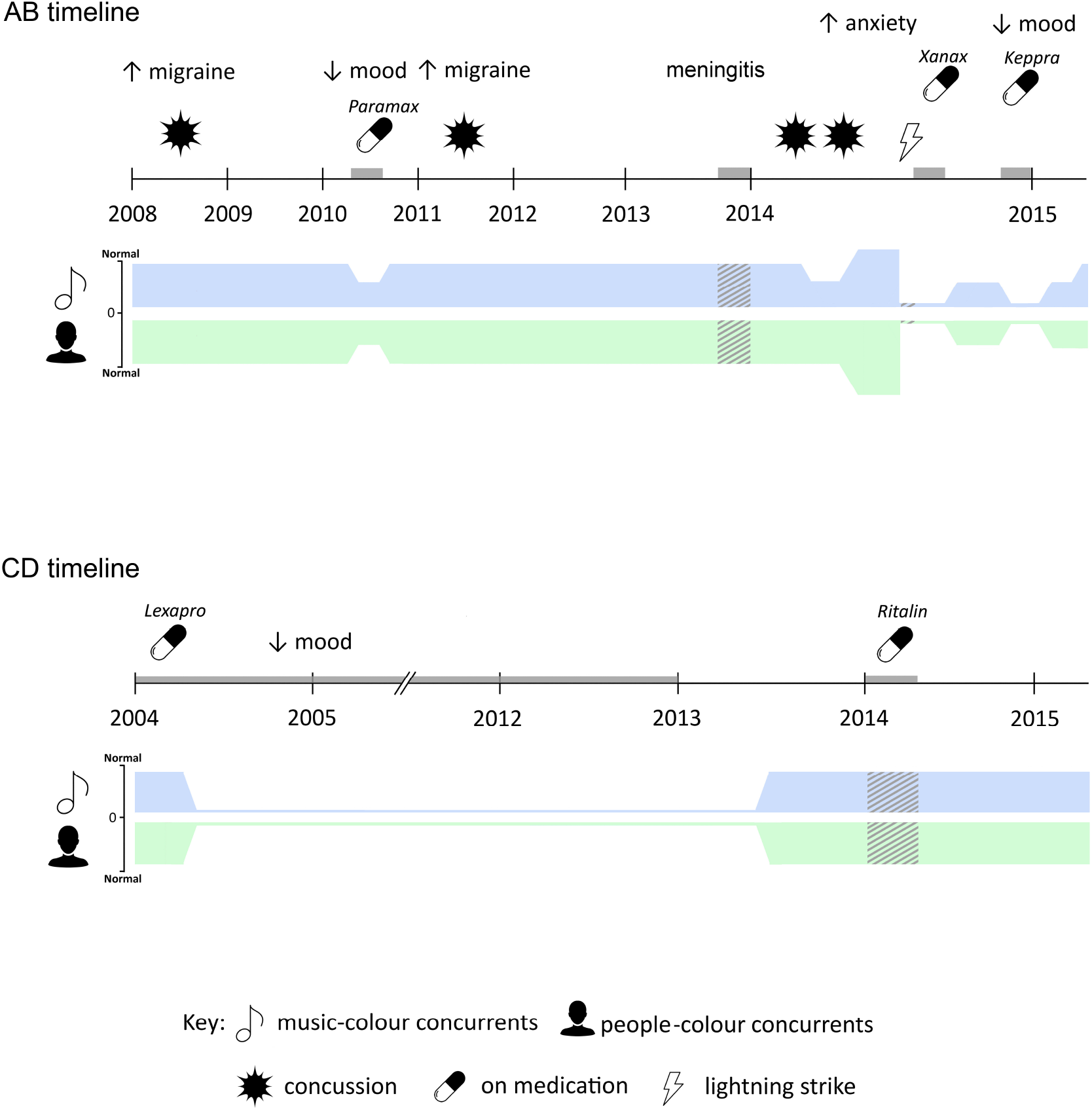
Timeline of events and resultant effects on synaesthetic experience for AB (upper) and CD (lower). Significant events depicted include concussions/head trauma, migraine, seizures, taking medication and lightning strike. Grey blocks indicate time spent on medication, or duration of meningitis. Hashed areas on colour bars indicate periods of altered synaesthetic colours. Icons were sourced from thenounproject.com and include Lightning Bolt by artworkbean, Pill by Sergey Demushkin, Burst by Bohdan Burmich, User by JM Waideaswaran and Music by Sherrinford.

**Table 1:** Data from the Online Synesthesia Battery (Eagleman et al., 2007) for AB (taken in 2012 and 2015) and CD (2015 only). For inducer-concurrent pairs, scores below 1.0 (in bold) indicate the presence of synaesthesia; for visual imagery, a Vividness of Visual Imagery Questionnaire (VVIQ) score above 3.0 indicated above average imagery. Negative scores on the projector-associator scale are indicative of associator-type.

|  | November 2012 | June 2015 |
| --- | --- | --- |
| **Subject AB** |  |  |
| *Battery Results* | Piano scale → colour: **0.57**<br>($< 1.0$ synaesthetic) | Piano scale → colour: 1.08 |
| | Instruments → colour: **0.78**<br>($< 1.0$ synaesthetic) | Instruments → colour: **0.45** |
| | Visual imagery: **4.72**<br>($>3$ higher than average) | Visual imagery: **5** |
| | Projector associator: **-1**<br>(negative = associator) | Chord → colour: 1.84<br>($< 1.0$ synaesthetic) |
| **Subject CD** |  |  |
| *Battery Results* | - | Instruments → colour: **0.57**<br>($< 1.0$ synaesthetic) |
| | - | Visual imagery: **4.22** ($>3$ higher than average) |
|  | - | Projector associator: **-0.33**<br>(negative = associator) |

By January 2015, AB had returned to normal cognitive and emotional functioning, but reports no restoration of her evoked colours at that time. Increased migraine in February caused heightened emotionality and distractibility, along with a partial return of her concurrents, albeit with altered or ‘wrong’ colours. Normal colour associations returned one month later. In June 2015, by which time she was seizure-free, AB completed a second Synesthesia Battery (see Table 1) and semi-structured interview with the authors. At this time, she scored positively for instrument-colour associations (0.45) and above average VVIQ (5.0), but not piano scale-colour (1.08) or chord-colour (1.84). AB's colour concurrents for people were also found to be highly consistent on a RGB colour-picker task, which included family members, friends and strangers (see Figure 3, Top).

**Figure 3:**
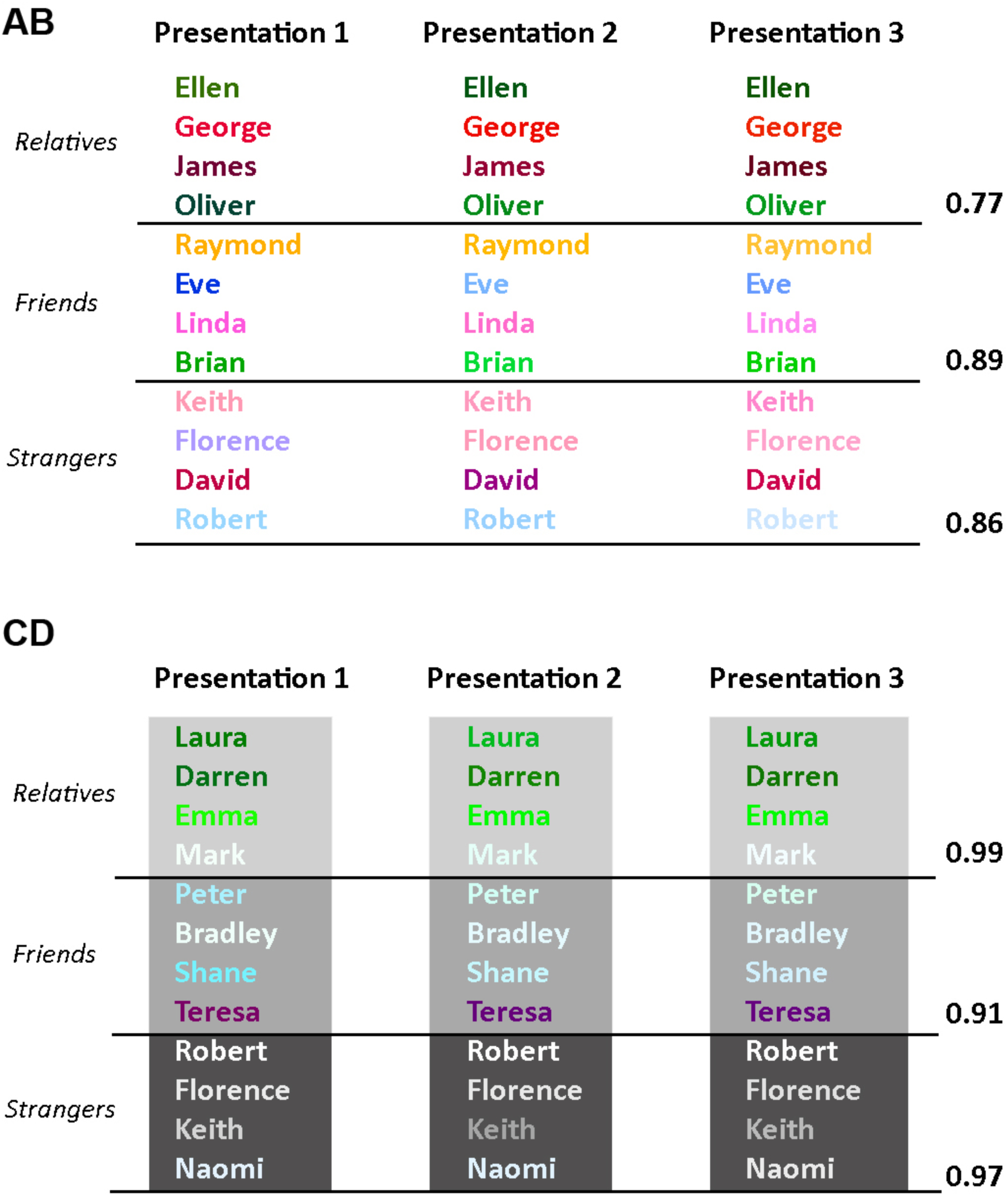
RGB-accurate colour concurrents for specific people for AB (upper) and CD (lower) on three successive occasions using a colour-picker. Mean correlation coefficients for red, green and blue values averaged across all three presentations are shown for each category: Relatives/Family Members; Friends; Strangers/Unknown. Names included in the Relatives and Friends categories were self-selected by AB and CD. Names in the Strangers/Unknown category were selected by the authors; these individuals were introduced briefly to AB and CD before the task. All names have been changed to preserve anonymity.

CD, a 31 year-old ambidextrous (right-hand dominant) male with a family history of autism, has experienced coloured auras around people and colours in response to music since childhood. Colours for people are automatically experienced around the person (usually the head) in the visual field, with specific colours associated with CD's mood and/or emotional reactions to the person (see Figure 1d). The degree to which auras are projected in space depends largely on the colour itself; for example, brown and black are typically perceived in sustained projections, while red appears much less frequently. When hearing music, he perceives semi-independent “colour blotches” in the centre and periphery of the visual field; unlike AB, these are entirely based on his emotional reactions to the music. He also has vague but consistent colour associations for some numbers, and visualises numeric functions in three-dimensional space; this can be used as an arithmetical aid. CD also possesses mild but consistent emotional associations for some letter groupings (e.g. ‘abc’ and ‘ijk’), as well as personification of letters (although these personalities tend to be inconsistent and were more prevalent during his childhood). Until 2015, CD viewed his synaesthetic experiences as being profoundly negative.

In 2004 (age 20), while taking Lexapro (*Escitalopram*) for Seasonal Affective Disorder, he first experienced a reduction of his visualisations, with a complete loss of colours for people and music occurring within 4-5 weeks. This absence persisted until the end of 2012 when he ceased taking the medication, resulting in the eventual return in summer 2013 of coloured concurrents to people and music (Figure 2). CD was briefly prescribed Ritalin (*Methylphenidate*) in 2014, leading to an alteration of his perceived colours for people; specifically, perceived colours were more purple-tinted and/or purple auras were enhanced. By July 2015, when he completed the Synesthesia Battery (see Table 1) and semi-structured interview, his original perceptual experiences (which he now views as positive) were returning and his visualisations were ‘becoming more coherent’. His battery scores indicated an associator-type synaesthete (-0.33) with instrument-colour associations (0.57) and above average VVIQ (4.22). CD's person-colour associations were found to be highly consistent across categories (see Figure 3, Bottom).

## Discussion

Both of these cases of music- and person-colour synaesthesia follow the classic descriptions of developmental synaesthesia, rather than being injury- or drug-induced. However, the synaesthetic experiences were modified by injury, by infection, migraine or seizure, or by a variety of drugs. Previous reports on the pharmacology of synaesthesia have focused mainly on the range of hallucinogenic drugs that can induce audiovisual synaesthesia-like experiences in non-synaesthetes or enhance them in synaesthetes, with a smaller number of reports of drugs that modulate the experience of developmental synaesthesia. Several authors have noted that many of these drugs, most notably the hallucinogens LSD, psilocybin and mescaline, act by modulating serotonin signalling, even leading to the proposal that mutations in serotonin pathway genes might underlie the condition (Brang & Ramachandran, 2008). Lexapro, which completely suppressed the synaesthetic experiences of CD for the eight years he was taking it, is a selective serotonin reuptake inhibitor, which is consistent with previous reported effects of fluoxetine (Brang & Ramachandran, 2008; Luke et al., 2012). However, the other drug effects reported here do not support such a selective relationship to serotonin signalling. Metoclopromide (present in Paramax) is an antagonist of dopamine receptors; Levetiracetam (Keppra) binds synaptic vesicle glycoprotein SV2A and modulates presynaptic L-type calcium channels, leading to reduced neurotransmitter release; Alprazolam (Xanax) is a benzodiazepine which allosterically increases chloride flux through GABA-A receptors, thus increasing inhibition; and Methylphenidate (Ritalin) is a norepinephrine and dopamine reuptake inhibitor. Thus, whatever the neural substrate of these synaesthetic experiences, they can clearly be modulated by a diverse range of drugs targeting very different pathways.

Remarkably, the drugs and other factors described here did not just change the intensity of the synaesthetic experience or affect whether it reached conscious awareness. They also led to qualitative changes in the nature of the concurrent percepts, specifically in the location in space of projected colours or in the specific colours induced. Furthermore, the changes were not short-lived – they persisted over days, weeks, months or years. However, despite the series of unfortunate events which caused successive alterations to AB's synaesthetic experiences over an eight-year period and the complete suppression of CD's synaesthesia also over an eight-year period, in both cases their synaesthesia has since returned to a similar level and qualitative nature as prior to these events. In the case of AB, direct comparison of colours for musical notes before and after the most severe set of events (recorded in 2012 and 2015; Figure 2a) shows a number of changes but an overall high level of consistency (0.69). These findings provide strong evidence that the neural substrates of synaesthetic associations, once they are consolidated in what is presumably an early critical period (Newell & Mitchell, 2015), remain “hard-wired” thereafter and can persist over very long periods even under conditions that alter or completely suppress the conscious synaesthetic experience itself.

## Acknowledgements

The authors gratefully acknowledge the contributions of Julia Januszweski and Laura Rai (Maynooth University) for transcribing the interviews with AB and CD, and Ríona McArdle (University of Newcastle) for the initial introduction to AB. Special thanks to AB and CD for their time, openness and willingness to engage with our tasks, and for feedback on the manuscript. Ethical approval for data collected was granted by the Maynooth University Research Ethics Committee, BSRESC-2015-001.

